# Network topology enables efficient response to environment in *Physarum polycephalum*

**DOI:** 10.1101/2022.11.09.515897

**Authors:** Siyu Chen, Karen Alim

## Abstract

The network-shaped body plan distinguishes the unicellular slime mould *Physarum polycephalum* in body architecture from other unicellular organisms. Yet, network-shaped body plans dominate branches of multi-cellular life such as in fungi. What survival advantage does a network structure provide when facing a dynamic environment with adverse conditions? Here, we probe how network topology impacts *P. polycephalum*’s avoidance response to an adverse blue light. We stimulate either an elongated, I-shaped amoeboid or a Y-shaped networked specimen and subsequently quantify the evacuation process of the light-exposed body part. The result shows that Y-shaped specimen complete the avoidance retraction in a comparable time frame, even slightly faster than I-shaped organisms, yet, at a lower almost negligible increase in migration velocity. Contraction amplitude driving mass motion is further only locally increased in Y-shaped specimen compared to I-shaped providing further evidence that Y-shaped’s avoidance reaction is energetically more efficient than in I-shaped amoeboid organisms. The difference in the retraction behaviour suggests that the complexity of network topology provides a key advantage when encountering adverse environments. Our findings could lead to a better understanding of the transition from unicellular to multicellularity.

## Introduction

Both mycelial fungi and plasmodial slime moulds such as *P. polycephalum* shape their body into extensive networks that are highly adaptive to their environment (1). With the interlaced network-shape *P. polycephalum* is at stark difference to the single cell amoeba state of its closest relative *Dictyostelium discoideum. D. discoideum* never transforms from the amoeba state but itself bridges into multicellular developmental stages by forming amoeba aggregates (2, 3). Instead *P. polycephalum* can transform its amoeba by fusion into a complex and dynamic multinucleate tubular network covering tens of centimeters (4). As a network-shaped multinucleate body plan evolved twice in mycelial fungi and plasmodial slime moulds, it suggests an evolutionary advantage of network over amoeboid shape.

*P. polycephalum* networks are highly dynamic, reorganizing when migrating and searching for food (5–7), responding to environmental conditions such as chemicals (8–10), humidity (11, 12), stiffness (13, 14), light conditions (11, 15–17) and temperature (18, 19), thereby distinguishing between attractive conditions such as nutrient-rich or adverse conditions such as blue light. When partly exposed to blue light the network rapidly reorganizes to evacuate from blue light (20, 21), as it harms the organism by inhibiting metabolism (22). By remodelling its network body in response to its environment *P. polycephalum* performs sophisticated, often termed ‘intelligent’ behaviours (20, 23, 24) like forming efficient adaptive and fault-tolerant networks connecting multiple food sources in a way comparable to optimized man-made transport networks (25, 26), or finding the shortest path through light (20). Being of network-shaped therefore allows for complex behaviours, but how do such advantages emerge when a tiny plasmodia grows beyond the amoeboid shape?

Amoeboid-shaped plasmodia have served as a desired model to investigate how migration arises from the coordination of the peristaltic contractions that enable mass flow (27– 31). Due to the reduced size, patterns of the acto-myosin driven cross-sectional contractions rhythmically constricting the single tube and thereby pumping the enclosed fluid volume forth and back can be quantified (29–31). Yet, how dynamics differ if the topology of elongated amoeboid, i.e. Ishaped plasmodium, becomes more complex by forming a first network node as in a Y-shaped network is unknown.

Here, we investigate the advantage of network topology by comparing the response of the simplest network state of a Y-shaped *P. polycephalum* plasmodium to the elongated Ishaped amoeboid plasmodia, in adverse conditions provided by localized blue light stimulating either a single branch of Yshaped or part of single-branched I shaped specimen. Quantifying in detail the dynamics of organism mass, motion and contraction dynamics of both Y-shaped and I-shaped side-by-side we discover that the higher complexity in topology of Y-shaped specimen allows them to evacuate from blue light more efficiently i.e. specimen move organism mass out of harmful blue light in a slightly shorter time period at less motion and lower overall contraction amplitude. Our finding therefore suggest that network topology may have a survival advantage compared to amoeboid shape when facing an adverse environment such as blue light.

## Methods

### Culturing of *P. polycephalum*

To ensure comparability of individual experiments in particular regarding the nutritional state, one day old microplasmodia grown in a liquid culture using the medium by Daniel and Rusch (32) with hematin (5 mgmL^*\**1^) instead of chicken embryo extract were used. The method to prepare plates with small I- and Y-shaped *P. polycephalum* plasmodia was adopted from the preparation of plasmodial networks as reported in (33). First, 1.5% agar plates were left to be air dried within the sterile environment with petri dish lids open for 5 min, before 1 mL of microplasmodia solution was added to the plates. The plates were then carefully tilted in all directions to ensure an even dispersion of plasmodia, this is to reduce the possibility of them fusing into one single plasmodium with complex network structure. After dispersion, the plates were left with lids opened to be air dried for another 15 min before being sealed with parafilm and incubated overnight under dark conditions at T = 25 °C. This second drying step is again crucial in preventing *P. polycephalum* from fusing together, it also ensures there are no visible traces of liquid medium during imaging which will affect the image quality. After plating, plates were used for imaging within 16-20 hours.

### Microscope setup and data acquisition

Images were recorded using a Zeiss Axio Zoom V16 with a Zeiss PlanNeofluar 1x/0.25 objective and a Hamamatsu ORCA-Flash 4.0 digital camera. The microscope setup is shielded from ambient light by a canopy, and a Lee 740 Aurora Borealis green filter was added in between the bright field light source and specimen to minimise the environmental stress on *P. polycephalum* (15).

After the petri dish was taken out from the incubator, it was sealed with parafilm and placed upside down onto the imaging stage to reduce the effect of agar drying out and visible condensation droplets during the experiment. The specimen was left under microscope light for 40 min before the start of the experiment to accommodate the change of *P. polycephalum* behaviour towards the change of the environment. The first 20 min of the experiment were carried out without blue light to capture a ‘baseline’ dynamics. Then, a blue light, created by a Zeiss HXP 200C light source at maximum intensity, and a Zeiss HE38 470/40 nm filter, was introduced at a predefined position. The intensity of the blue light used varies depending on the optical zoom, in the following experiments it was between 150 and 250 Wm^*\**2^ as was measured by a Thorlabs PM16-122 Power Meter at the sample stage.

The Zeiss Zen 2 (Blue Edition) software was used for imaging. An image was acquired every 6 s.

The position of the blue light need to be predefined in Zen software before the start of the experiments, meanwhile the *P. polycephalum* could have migrated away from the blue light region during the first hour before the blue light was triggered. In those cases, the experiment was aborted, and a new petri dish with a sample of the right topology was taken from the incubator, to ensure all samples were in a similar state when meeting the blue light.

### Image analysis

The resulting experimental images were first passed through a custom-written MATLAB code following the steps described in (33) to extract a measure of organism volume and its dynamics in terms of the intensities of transmitted light at every pixel along the backbone of the specimen. Briefly, during this process, the first step was to remove the optical defects (e.g. vignetting and uneven lighting) of each image with the rolling-ball method. With this background-adjusted-image, a threshold was applied to separate the network from the background. For the extracted binary network, there might be small holes in the structure and rough edges due to the thresholding, thus, small holes were filled and single-pixel edges were smoothed before the biggest structure was extracted as a binary ‘mask’. This binary ‘mask’ was used to extract the pixel information of the *P. polycephalum* network from the background-adjusted-image, which was later used to calculate the mass and the weighted-centroid.

In the next step, an iterative custom written etching process was applied to the mask to extract a ‘skeleton’ of the network. For each pixel on the network, a local diameter, which was calculated as the largest fitting disk radius around the point within the mask, was calculated. Within this disk, the average intensity was computed and saved with the skeleton data. Because the diameter has a lower resolution than the intensity, the intensity data was used for the calculation of amplitude and frequency.

As *P. polycephalum* performs avoidance reaction and migrates out of the blue light region, a reference skeleton image that captures all organism location data throughout the experiment was also created in order to achieve a kymograph-like file (‘intensity kymograph’) where the time evolution of the network pixel intensity along the tube can be recorded and tracked. This reference image was achieved by first overlay and then etch all skeleton data. Next, for each time frame, each pixel on the instantaneous skeleton data was mapped on to a corresponding pixel in the reference image using the shortest distance as criteria. Lastly, the corresponding intensity data at each time frame for each pixel can be loaded onto the reference position, and the kymograph was generated.

## Results

### I- and Y-shapes require similar time to evacuate from a blue light region

To study how network topology impacts the response of *P. polycephalum* to blue light stimuli, we collected time series of bright-field images of the organism’s evacuation response over approximately 2 h, during which a single, localised blue light stimulus was applied on a part of the organism (Fig. 1A). For I-shaped specimen, individual experiments differ in the fraction of the elongated organism that was exposed to light. To address which advantage an additional branch provide for the retraction of a ‘branch’ from harmful light, we only expose one out of the three ‘branches’ connected into a Y for the Y-shaped specimen to blue light.

**Fig. 1.**
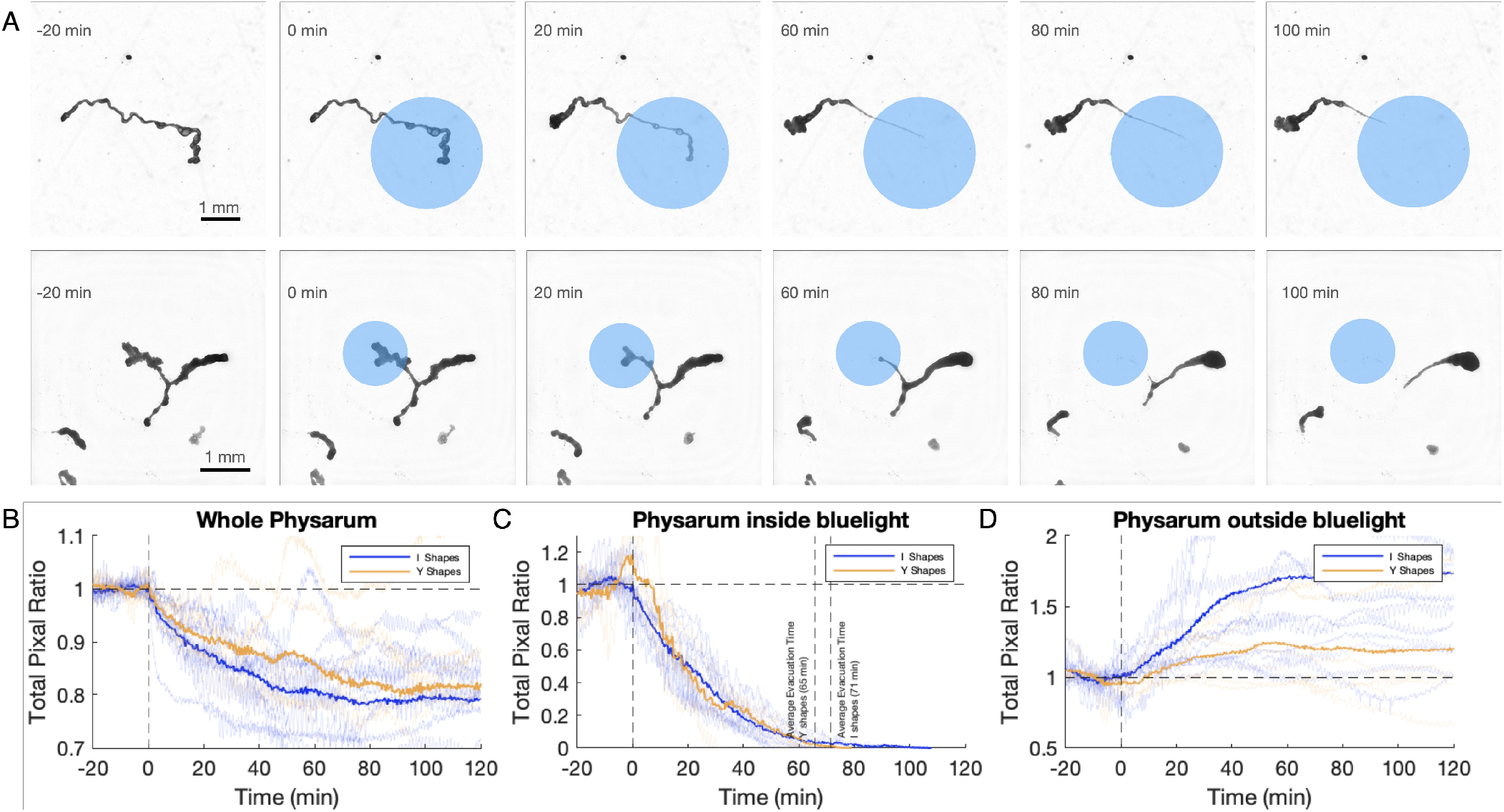
Y-shaped *P. polycephalum* evades blue light region slightly faster than I-shaped, while I-shaped suffer on average a slightly larger loss in overall mass. (A) Sequential bright-field images of a previously stable *P. polycephalum* evading the localized blue light stimulus (blue circle) applied at 0 min. (B) Averaged change of *P. polycephalum* total mass for the whole organism shows a statistically stronger decrease in mass for I-shaped specimen. (C) Averaged change of the *P. polycephalum* mass for the specimen part exposed to blue light indicates a statistically slightly faster evacuation for Y-shaped specimen. The blue and yellow dashed line indicates average evacuation time for I- and Y-shaped organisms. Respective dotted line indicates average evacuation time for specimen subrgroup with less than half of the specimen exposed to light initially. (D) Evacuation response is compensated by averaged change of *P. polycephalum* specimen mass outside of blue light stimulus. The y-axis presents the ratio of the instantaneous mass (total pixel value) over the baseline value (mean pixel value before blue light was applied), respectively. Bold yellow line indicates averaged I-shape data (N = 12) and bold blue line averaged Y-shape data (N = 6). Light blue and yellow lines in the background are the change in the total mass for individual experiments.

During the 20 min before the application of blue light, where the organism is freely living on the agar plate (the ‘baseline’), the body mass, approximated by the sum of recorded bright-field intensity values related to the organisms mass by Lambert-Beer law, is conserved (Fig. 1B). As the organism is continuously consuming energy (34) but here is followed in non-nutritious environment we expect specimen mass to decrease over time (35).

For each experiment, the instantaneous body mass was compared to the average of the baseline value, by taking the ratio of the two masses. We choose to take the ratio instead of difference measure as the ratio allows for ‘non-dimensional’ comparison of specimen differing in initial statistics and is more robust against fluctuations in body mass arising as body mass diminishes particularly in the light-exposed region. To obtain generalised I-shaped and Y-shaped behaviour, we then averaged the data points for I- and Y-shapes, respectively. As shown in Fig. 1B, the whole *P. polycephalum* mass decreases under the exposure to blue light. Change in mass is significantly larger than mass fluctuations triggered by the exposure to microscopy light between 60 min to 20 min before onset of light exposure, see Supplemental Fig. 1. Mass decrease is slightly stronger for I-shaped than for Y-shaped specimen. To quantify the evacuation response we then followed the *P. polycephalum* mass exposed to blue light over time (Fig. 1C). On average I-shaped took 71 min (N = 12, SD = 17) and Y-shaped took 65 min (N = 6, SD = 18) to evacuate as indicated by the dashed lines in Fig. 1C. While for extreme cases, the slowest Y-shape took 79 min to complete the evacuation, the slowest I-shape took 108 min to complete evacuation. The slightly faster evacuation of Y versus I-shaped also holds when grouping data by the mass fraction exposed to light. If less than half of the organism was illuminated by blue light upon onset of exposure, I-shaped *P. polycephalum* took 66 min (N = 6, SD = 16) while the Y-shaped organism took 63 min (N = 5, SD = 18) for evacuation, see dotted lines in Fig. 1C. The evacuation from the region of blue light is accompanied by the specimen’s part outside of blue light increasing over time (Fig. 1D). Note, that as total pixel ratios are normalized individually for panels B, C and D in Fig. 1 data inside and outside blue light only add up to the whole *P. polycephalum*data when weighted with their respective normalization.

Despite only subtle differences in the overall mass shrinkage and evacuation time, the process of how evacuation is achieved differs strongly between I- and Y-shaped specimen. For I-shaped, the part inside the blue light undergoes both thinning and retracting process (see i.e. Fig. 1A at 60 min and Supplemental Videos), while Y-shaped only smoothly retract the light exposed part.

### I-shapes exhibit higher velocity during evacuating from the blue light region

To measure the differences in the evacuation process we next quantify the changes in motion velocity of the whole organisms as well as the individual parts exposed or not exposed to blue light. Here, the instantaneous velocity is calculated from the position change of the weighted centroid of the *P. polycephalum* between frames. Again as a baseline velocity we calculated the average velocity during the 20 min before blue light was applied to individual *P. polycephalum*, and the instantaneous velocities were compared to this baseline velocity of the whole or the specimen part outside or inside blue light, respectively. Note, that the non-circular specimen shape is limiting the accuracy of centroid motion representing specimen motion. To assess long-term *P. polycephalum* motion on the scales of hours tracking the position of growth fronts is more advisable (36). Yet, to assess, here, the speed of the evacuation process the centroid motion is a better suited precisely because of the complex shape changes. As blue light exposure has been implicated to change the mechanical properties of both *P. polycephalum* tube wall and cytoplasm (21), we suspect these mechanical changes can have a direct implication on the velocity. Hence, the velocity data of individual specimen was binned according to the ratio of specimen’s body mass exposed to blue light compared to that of the whole *P. polycephalum* body. Different specimen have different proportions of their body initially exposed to blue light, with particularly Y-shaped being exposed less than 60%, but as evacuation continues this proportion always shrinks to zero. Thus, the x-axis in Fig. 2 can be read as inverse time.

**Fig. 2.**
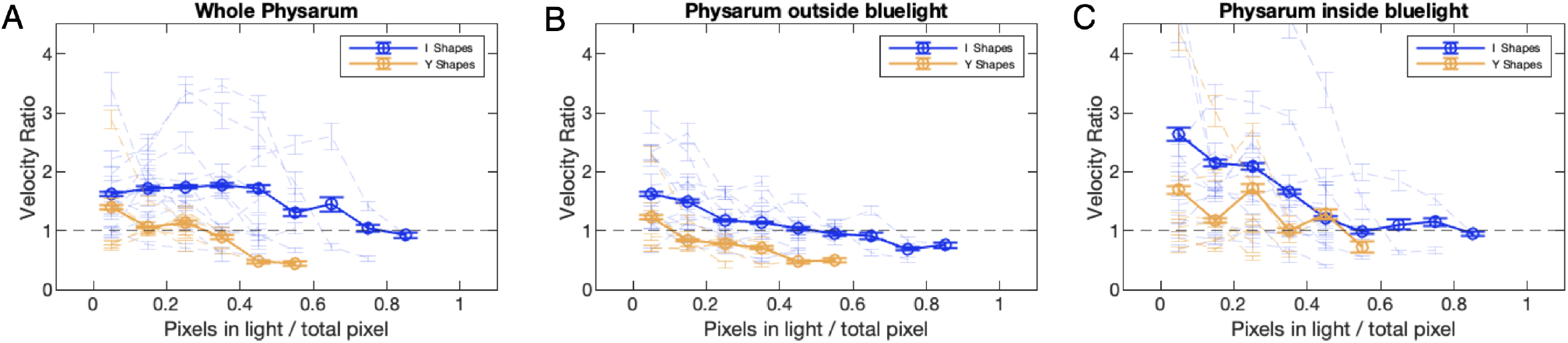
Change of velocity bigger for I- than Y-shaped *P. polycephalum*. Ratio of velocity (A) for the whole specimen body, (B) for the specimen part outside of blue light, and (C) for the specimen part under blue light, to their averaged velocities within 20 min prior to light exposure, respectively. Velocities ratios were binned into the relative mass of *P. polycephalum* in blue light relative to the whole body. Error bars denote standard errors of data. For both topologies, there is a clear increase of velocity for the region directly under blue light. Meanwhile, the response from the region outside of the blue light differs, which resulted in the overall higher evacuation velocity for the I-shapes. Each light blue and yellow line in the background represents a single data set, and bold blue and yellow lines represent the ensemble average within each bin.

**Fig. 3.**
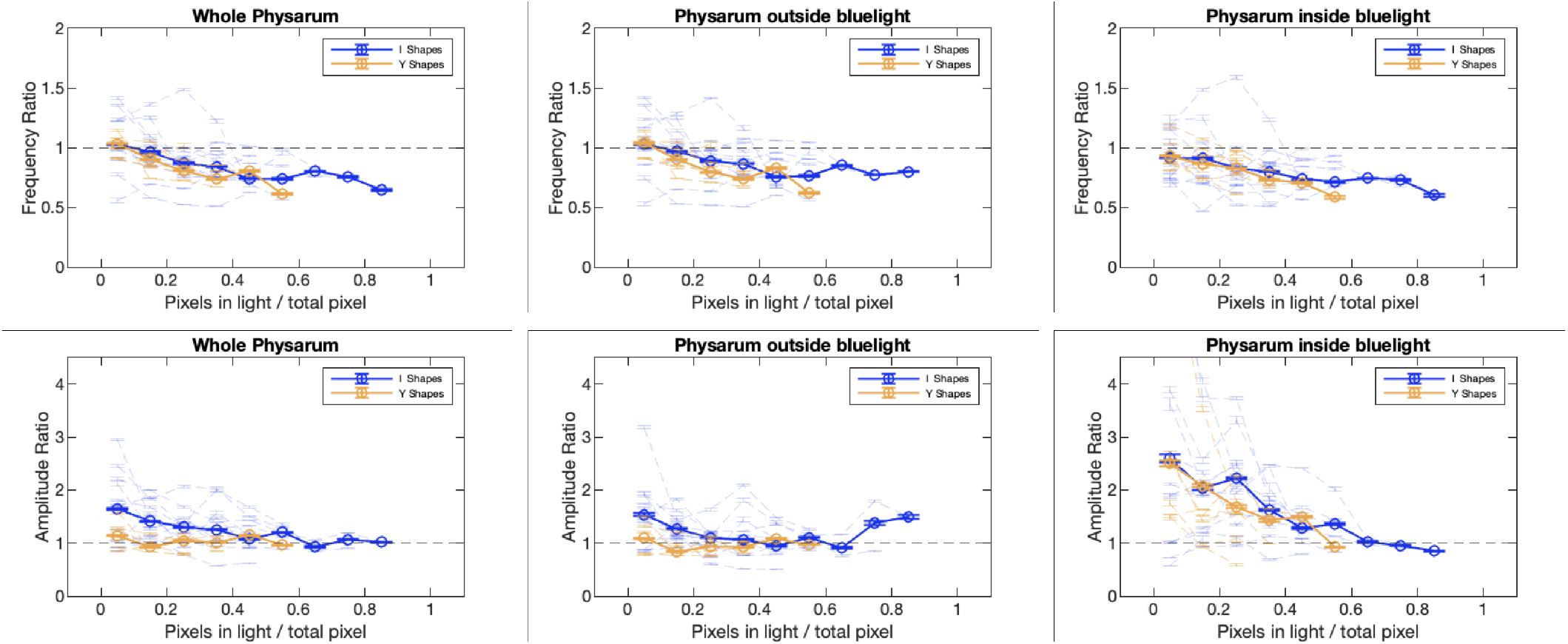
Contraction frequency equally decreases upon light exposure whereas contraction amplitude increases more pronounced for I-shaped specimen. Ratio of frequency (upper row) and amplitude (lower row) to base frequency and amplitude within the 20 min prior to light exposure, respectively, for I- and Y-shaped *P. polycephalum* versus area percentage of instantaneous light exposure as in Fig. 2. Error bars denote standard error. For both topologies there is an equal decrease in contraction frequency upon light exposure and recovery upon evacuation. The contraction amplitude, on the contrary, exhibits an increase in the organism part under blue light for both topologies, but I-shaped also experience an increase in the organism part outside of blue light.

First, we focus on the overall dynamics, and find that both I- and Y-shaped *P. polycephalum* increase their velocity during the evacuation process continues (Fig. 2A). On average, for the I-shaped, the velocity increases on by 52%, exceeding the coefficient of variation of 27% obtained from the 20 min time window prior to light exposure. Similarly for the Y-shaped, the velocity increases on average by 27%, exceeding its prior to light exposure coefficient of variation of 19%. Notably, the velocity of Y-shaped is always lower than that of I-shaped. After being exposed to blue light I-shaped first continue the overall motion at their baseline velocity, only as the evacuation process continues the velocity increases. Velocity dynamics reach a plateau as the organism is halfway out of the blue light, maxing out at 1.7 times the baseline velocity. For Y-shaped, instead of continuing with the baseline velocity, it lowers its locomotion velocity upon the onset of blue light. Over time as it evacuates, it slowly recovers to the baseline velocity. It only uses a higher overall velocity at the last step of the evacuation process when there is less than 10% of the mass remaining in the blue light.

Distinguishing *P. polycephalum* into a specimen’s part outside of blue light and a part exposed to blue light, the weighted centroids of each part and, therefrom the velocities were recalculated for each part individually (Fig. 2B, C). For I-shaped organism, both parts show a constant increase in the instantaneous velocity compared to their individual baselines as evacuation progresses. Yet, the increase in velocity outside of blue light is much less vigorous as for the exposed part. This is partially due to the nature of an evacuation process, where in the later stage, for the *P. polycephalum* inside blue light, a small change in the mass can have a large impact in shifting the weighted centroid and thus result in a large velocity. This gradual decrease of the mass fraction inside the blue light also means that the contribution of the velocity increase from this light exposed part to the overall specimen velocity is constantly decreasing, which explains why the velocity of the whole I-shaped organism reaches a plateau while both velocities inside and outside the blue light increase.

For Y-shaped, the dynamics of the organism part outside of blue light resembles the overall velocity, recovering steadily from a decrease in motion velocity upon the impact of blue light. For the part of the organism under blue light, an increase in motion velocity similar but not as steady and fast as for I-shaped organisms is observed. Interestingly, there is an oscillation pattern for the inside the blue light region. From the data recordings, it seems that the organism deploys alternating dynamics during the evacuation process. Namely, high velocity where mass is transported out of the branch affected by blue light, and low velocity where the non-exposed branches outside of the blue light redistribute the newly acquired excess mass.

From the slightly faster evacuation of Y-shaped versus Ishaped observed in the previous section, here, finding motion velocity of I-shaped to be consistently faster than Y-shaped is against our intuition if we assume that mass motion is driving evacuation. Thus, there needs to be a more subtle mechanism to evacuate the light-exposed organism part that gives advantage to more complex Y-shaped network topologies over I-shaped amoeboid topologies. Due to their simple topology, I-shaped *P. polycephalum* are constrained in how their body mass can be reallocated using shuttle streaming. Body mass that needs to be moved out of blue light either needs to be pumped by the light exposed part or sucked by the non-exposed part by shuttle flow into the part outside of blue light. In the Y-shaped topology, however, the two branches outside of blue light do not need to undergo a net motion as they may symmetrically inflate to take up mass. To identify the pumping dynamics we next turn to quantify the contraction dynamics.

### A. Blue light triggers increase in amplitude in light exposed part but also non-exposed part for I-shaped

*P. polycephalum* uses peristaltic pumping as a transport mechanism (37), we, hence, analysed the contraction characteristics, namely the instantaneous frequency and amplitude, for I- and Y-shaped organisms. Both frequency and amplitude are extracted individually on the pixel level from pixel intensity time dynamics (‘intensity kymograph’) and then subsequently averaging over pixels in respective parts of an organism. To achieve this, the time-evolution of the intensity data was first ‘detrended’ with a moving-average filter and smoothed with a Gaussian filter, this is to further eliminate the difference in energy level of *P. polycephalum* that may exist and could induce bias. Then the Hilbert transform was performed to obtain an ‘analytic signal’. From which, the amplitude can be easily obtained by taking the absolute value, and the frequency can be obtained by calculating the phase angle and then unwrapping it. More details regarding the Hilbert transform can be found in (33).

Similar to Fig. 2, frequency and amplitude change are quantified relative to the baseline, the average frequency and amplitude during the 20 min prior to light onset, and subsequently binned according to the relative mass exposed to light. The changes in frequency of both I- and Y-shaped almost collapse regardless of which part of the organism is considered. Upon blue light exposure frequency initially decreases (15, 38–40) to then slowly recover to the baseline frequency as the evacuation process continues.

On the contrary, upon light exposure the amplitude increases both for I-shaped and Y-shaped organisms within the organism part exposed to blue light, also see Fig. S1 for Y-shaped individuals. However, for the part outside of blue light, the two topologies showed different responses: for Y-g shaped plasmodia the amplitude was maintained at around the baseline value, while for I-shaped plasmodia an increase in amplitude was observed as the evacuation proceeded. The differences in the dynamics in organism’s part outside of blue light dominate the overall *P. polycephalum* body amplitude dynamics. Note, that here as in Fig. 2 the individual normalization of the specimen parts outside and inside blue light results in the whole organism dynamics being a non-linear function of the respective parts.

In a peristaltic wave, as frequency decreases, pumping performance decreases, yet, an increase in contraction amplitude counteracts. When the amplitude increase outperforms the frequency decrease, overall the pumping performance may increase. Here, as amplitude increase rises to more than twofold, at a less than half-fold decrease in frequency, the exposure to blue light seems to up-regulate pumping performance in the light-exposed part. The localized up-regulation of pumping performance appears to be sufficient to efficiently evacuate the light-exposed part for Y-shaped specimen. For the simpler topology of I-shaped amoeboid organisms contraction amplitude also increases in the non-exposed body parts. This more wide-spread increase in contraction amplitude suggests that a similar effectiveness in evacuation speed for I-shaped specimen can only be achieved if also the pumping i.e. sucking performance in the non-exposed organism part is increased. However, up-regulating contraction amplitude in a larger portion of an organism is likely to be more energetically costly (41) and, thus, may account for the observation of a stronger mass consumption of I-shaped specimen during evacuation than Y-shaped (Fig. 1B). The more complex albeit still simple topology of Y-shaped organism body may therefore allow for a more efficient response to adverse conditions.

## Discussion

Comparing the dynamics of the response to localized blue light exposure of elongated amoeboid I-shaped versus network-state Y-shaped *P. polycephalum*, we find that Y-shaped are slightly faster in evacuating their light exposed body part than I-shaped. All the while I-shaped body parts exhibit an overall faster motion velocity and increase contraction amplitude across the entire organism while in Y-shaped only the light exposed body part increases contraction amplitude. This increased dynamics of I-shaped go at the expense of their body mass decreasing statistically slightly stronger than Y-shaped. We therefore conclude that the network topology represented by the Y-shaped organisms requires less energy and is, thus, more efficient in evacuating body parts exposed to harmful conditions.

The decisive difference in the dynamics of contractions channelling the mass movement is the across-organism increase in contraction amplitude in I-shaped versus the localized contraction amplitude increase in Y-shaped specimen. It is likely that this difference is deeply rooted in the contrasted topologies. Within amoeboid I-shaped organisms dynamics along the organism are inherently coupled and an overall increase in pumping efficiency is best to quickly increase the mass movement along the peristaltic pressure drop along the organism (42). For Y-shaped networks, however, the two unexposed branches may maintain a low pressure at the network node and therefore passively facilitate the mass motion ignited by the increase in pumping of the light exposed network arm. This could also explain the seemingly two different escaping mechanisms we are observing, namely the I-shaped needs as much body length as possible to generate the maximum amplitude, hence, choose to decouple the thinning and retracting process during evacuation, while Y-shaped organisms can benefit from the coordination of the two branches outside the blue light and, hence, smoothly retract the light exposed part without a visible thinning step. It would be, here, instructive to built on previous work (10, 21, 43–45) to mathematically model how potential mechanical changes to the actin cortex may lead to the observed differences in amplitude and frequency changes in the Iversus Y-shaped morphologies.

Network complexity is multiplexed. It ranges from the information storage opening up from the pattern of thicker and thinner tubes (46, 47), to dynamic stability arising from symmetries in networks topology (48). Here, we demonstrated, the mere existence of a network node can allow for behavioural advantage. The puzzle of how networked body forms emerged gets added a new puzzle piece that may also shine light on the evolutionary process toward multicellular organisms and the beginning of the more complicated forms of life.

## ACKNOWLEDGEMENTS

We sincerely thank Jean-Daniel Julien for insightful discussions. This project has received funding from the Max Planck Society and the European Research Council (ERC) under the European Union’s Horizon 2020 research and innovation programme (grant agreement No. 947630, FlowMem) to K.A..

## Bibliography

1. Mark D. Fricker, Lynne Boddy, Toshiyuki Nakagaki, and Daniel P. Bebber. Adaptive net-works, theory, models and applications. Understanding complex systems, pages 51–70, 2009.

2. L Eichinger, JA Pachebat, G Glöckner, M-A Rajandream, R Sucgang, M Berriman, J Song, R Olsen, Ku Szafranski, Q Xu, et al. The genome of the social amoeba dictyostelium discoideum. Nature, 435(7038):43–57, 2005.

3. Christian Westendorf, Jose Negrete, Albert J Bae, Rabea Sandmann, Eberhard Bodenschatz, and Carsten Beta. Actin cytoskeleton of chemotactic amoebae operates close to the onset of oscillations. Proceedings of the national academy of sciences, 110(10):3853–3858, 2013.

4. Helmut Sauer and Peter W Barlow. Developmental biology of Physarum, volume 11. CUP Archive, 1982.

5. NB Matveeva, SI Beilina, and VA Teplov. The role of phosphoinositide-3-kinase in the control of shape and directional movement of the physarum polycephalum plasmodium. Biophysics, 53(6):533–538, 2008.

6. Chris R Reid, Madeleine Beekman, Tanya Latty, and Audrey Dussutour. Amoeboid organism uses extracellular secretions to make smart foraging decisions. Behavioral ecology, 24 (4):812–818, 2013.

7. Daniel Schenz, Yasuaki Shima, Shigeru Kuroda, Toshiyuki Nakagaki, and Kei-Ichi Ueda. A mathematical model for adaptive vein formation during exploratory migration of physarum polycephalum: routing while scouting. Journal of physics D: applied physics, 50(43): 434001, 2017.

8. Randall L Kincaid and Tag E Mansour. Measurement of chemotaxis in the slime mold physarum polycephalum. Experimental cell research, 116(2):365–375, 1978.

9. J Kukulies, W Stockem, and KE Wohlfarth-Bottermann. Caffeine-induced surface blebbing and budding in the acellular slime mold physarum polycephalum. Zeitschrift für naturforschung C, 38(7-8):589–599, 1983.

10. M Radszuweit, H Engel, and M Bär. A model for oscillations and pattern formation in protoplasmic droplets of physarum polycephalum. The european physical journal special topics, 191(1):159–172, 2010.

11. Leokadia Rakoczy. The myxomycete physarum nudum as a model organism for photobiological studies. Berichte der deutschen botanischen gesellschaft, 86(1-4):141–164, 1973.

12. K Takahashi, A Takamatsu, Z-S Hu, and Y Tsuchiya. Asymmetry in the self-sustained oscillation ofphysarum plasmodial strands. Protoplasma, 197(1):132–135, 1997.

13. Atsuko Takamatsu, Eri Takaba, and Ginjiro Takizawa. Environment-dependent morphology in plasmodium of true slime mold physarum polycephalum and a network growth model. Journal of theoretical biology, 256(1):29–44, 2009.

14. Nirosha J Murugan, Daniel H Kaltman, Paul H Jin, Melanie Chien, Ramses Martinez, Cuong Q Nguyen, Anna Kane, Richard Novak, Donald E Ingber, and Michael Levin. Mechanosensation mediates long-range spatial decision-making in an aneural organism. Advanced materials, 33(34):2008161, 2021.

15. Masakatsu Hato, Tetsuo Ueda, Kenzo Kurihara, and Yonosuke Kobatake. Phototaxis in true slime mold physarum polycephalum. Cell structure and function, 1(3):269–278, 1976.

16. Toshiyuki Nakagaki, Shoji Umemura, Yasutaka Kakiuchi, and Tetsuo Ueda. Action spectrum for sporulation and photoavoidance in the plasmodium of physarum polycephalum, as modified differentially by temperature and starvation. Photochemistry and photobiology, 64 (5):859–862, 1996.

17. B Rodiek and MJB Hauser. Migratory behaviour of physarum polycephalum microplasmodia. The european physical journal special topics, 224(7):1199–1214, 2015.

18. KE Wohlfarth-Bottermann. Oscillating contractions in protoplasmic strands of physarum: Simultaneous tensiometry of longitudinal and radial rhythms, periodicity analysis and temperature dependence. Journal of experimental biology, 67(1):49–59, 1977.

19. Z Hejnowicz and KE Wohlfarth-Bottermann. Propagated waves induced by gradients of physiological factors within plasmodia ofphysarum polycephalum. Planta, 150(2):144–152, 1980.

20. Toshiyuki Nakagaki, Makoto Iima, Tetsuo Ueda, Yasumasa Nishiura, Tetsu Saigusa, Atsushi Tero, Ryo Kobayashi, and Kenneth Showalter. Minimum-risk path finding by an adaptive amoebal network. Physical review letters, 99(6):068104, 2007.

21. Felix K Bäuerle, Stefan Karpitschka, and Karen Alim. Living system adapts harmonics of peristaltic wave for cost-efficient optimization of pumping performance. Physical review letters, 124(9):098102, 2020.

22. Thomas Schreckenbach, Barbl Walckhoff, and Cornelia Verfuerth. Blue-light receptor in a white mutant of physarum polycephalum mediates inhibition of spherulation and regulation of glucose metabolism. Proceedings of the national academy of sciences, 78(2):1009–1013, 1981.

23. Toshiyuki Nakagaki and Robert D Guy. Intelligent behaviors of amoeboid movement based on complex dynamics of soft matter. Soft matter, 4(1):57–67, 2008.

24. Audrey Dussutour, Tanya Latty, Madeleine Beekman, and Stephen J Simpson. Amoeboid organism solves complex nutritional challenges. Proceedings of the national academy of sciences, 107(10):4607–4611, 2010.

25. Toshiyuki Nakagaki, Hiroyasu Yamada, and Ágota Tóth. Maze-solving by an amoeboid organism. Nature, 407(6803):470–470, 2000.

26. Atsushi Tero, Seiji Takagi, Tetsu Saigusa, Kentaro Ito, Dan P Bebber, Mark D Fricker, Kenji Yumiki, Ryo Kobayashi, and Toshiyuki Nakagaki. Rules for biologically inspired adaptive network design. Science, 327(5964):439–442, 2010.

27. Shun Zhang, Robert D Guy, Juan C Lasheras, and Juan C Del Álamo. Self-organized mechano-chemical dynamics in amoeboid locomotion of physarum fragments. Journal of physics D: Applied physics, 50(20):204004, 2017.

28. Owen L Lewis, Shun Zhang, Robert D Guy, and Juan C Del Alamo. Coordination of contractility, adhesion and flow in migrating physarum amoebae. Journal of the royal society interface, 12(106):20141359, 2015.

29. Owen L Lewis and Robert D Guy. Analysis of peristaltic waves and their role in migrating physarum plasmodia. Journal of physics D: applied physics, 50(28):284001, 2017.

30. Philipp Fleig, Mirna Kramar, Michael Wilczek, and Karen Alim. Emergence of behaviour in a self-organized living matter network. Elife, 11:e62863, 2022.

31. Beatrice Rodiek, Seiji Takagi, Tetsuo Ueda, Marcus Hauser, et al. Patterns of cell thickness oscillations during directional migration of physarum polycephalum. European biophysics journal, 44(5):349–358, 2015.

32. John W Daniel and Harold P Rusch. The pure culture of physarum polycephalum on a partially defined soluble medium. Microbiology, 25(1):47–59, 1961.

33. Felix K Bäuerle, Mirna Kramar, and Karen Alim. Spatial mapping reveals multi-step pattern of wound healing in physarum polycephalum. Journal of physics D: applied physics, 50(43): 434005, 2017.

34. Atsuko Takamatsu, Takuma Gomi, Tatsuya Endo, Tomo Hirai, and Takato Sasaki. Energy-saving with low dimensional network in Physarum plasmodium. Journal of physics D: Applied physics, 50(15):154003, 2017.

35. Sophie Marbach, Karen Alim, Natalie Andrew, Anne Pringle, and Michael P. Brenner. Pruning to Increase Taylor Dispersion in Physarum polycephalum Networks. Physical review letters, 117(17):178103, 2015.

36. Shigeru Kuroda, Seiji Takagi, Toshiyuki Nakagaki, and Tetsuo Ueda. Allometry in Physarum plasmodium during free locomotion: size versus shape, speed and rhythm. Journal of experimental biology, 218(23):3729–3738, 2015. ISSN 0022-0949.

37. Karen Alim, Gabriel Amselem, François Peaudecerf, Michael P Brenner, and Anne Pringle. Random network peristalsis in physarum polycephalum organizes fluid flows across an individual. Proceedings of the national academy of sciences, 110(33):13306–13311, 2013.

38. Z Baranowski, Z Shraideh, and KE Wohlfarth-Bottermann. Which phase of the contraction-relaxation cycle of cytoplasmic actomyosin in physarum is modulated by blue light? Cell biology international reports, 6(9):859–865, 1982.

39. I Block and KE Wohlfarth-Bottermann. Blue light as a medium to influence oscillatory contraction frequency in physarum. Cell biology international reports, 5(1):73–81, 1981.

40. Atsuko Takamatsu, Kengo Takahashi, Makoto Nagao, and Yoshimi Tsuchiya. Frequency coupling model for dynamics of responses to stimuli in plasmodium of physarum poly-cephalum. Journal of the physical society of Japan, 66(6):1638–1646, 1997.

41. Christina Oettmeier, Klaudia Brix, and Hans-Günther Döbereiner. Physarum poly-cephalum—a new take on a classic model system. Journal of physics D: applied physics, 50(41):413001, 2017.

42. Meijing Li and James G Brasseur. Non-steady peristaltic transport in finite-length tubes. Journal of fluid mechanics, 248:129–151, 1993.

43. Thomas P Stossel. Contribution of actin to the structure of the cytoplasmic matrix. The journal of cell biology, 99(1 Pt 2):15s, 1984.

44. D Takagi and NJ Balmforth. Peristaltic pumping of viscous fluid in an elastic tube. Journal of fluid mechanics, 672:196–218, 2011.

45. Jean-Daniel Julien and Karen Alim. Oscillatory fluid flow drives scaling of contraction wave with system size. Proceedings of the national academy of sciences, 115(42):10612–10617, 2018.

46. Mirna Kramar and Karen Alim. Encoding memory in tube diameter hierarchy of living flow network. Proceedings of the national academy of sciences, 118(10):e2007815118, 2021.

47. Komal Bhattacharyya, David Zwicker, and Karen Alim. Memory Formation in Adaptive Networks. Physical review letters, 129(2):028101, 2022.

48. Francis G. Woodhouse, Aden Forrow, Joanna B. Fawcett, and Jörn Dunkel. Stochastic cycle selection in active flow networks. Proceedings of the national academy of sciences, 113(29):8200–8205, 2016.

