## Supplementary material for "Network topology enables efficient response to environment in *Physarum polycephalum*": Figure S2

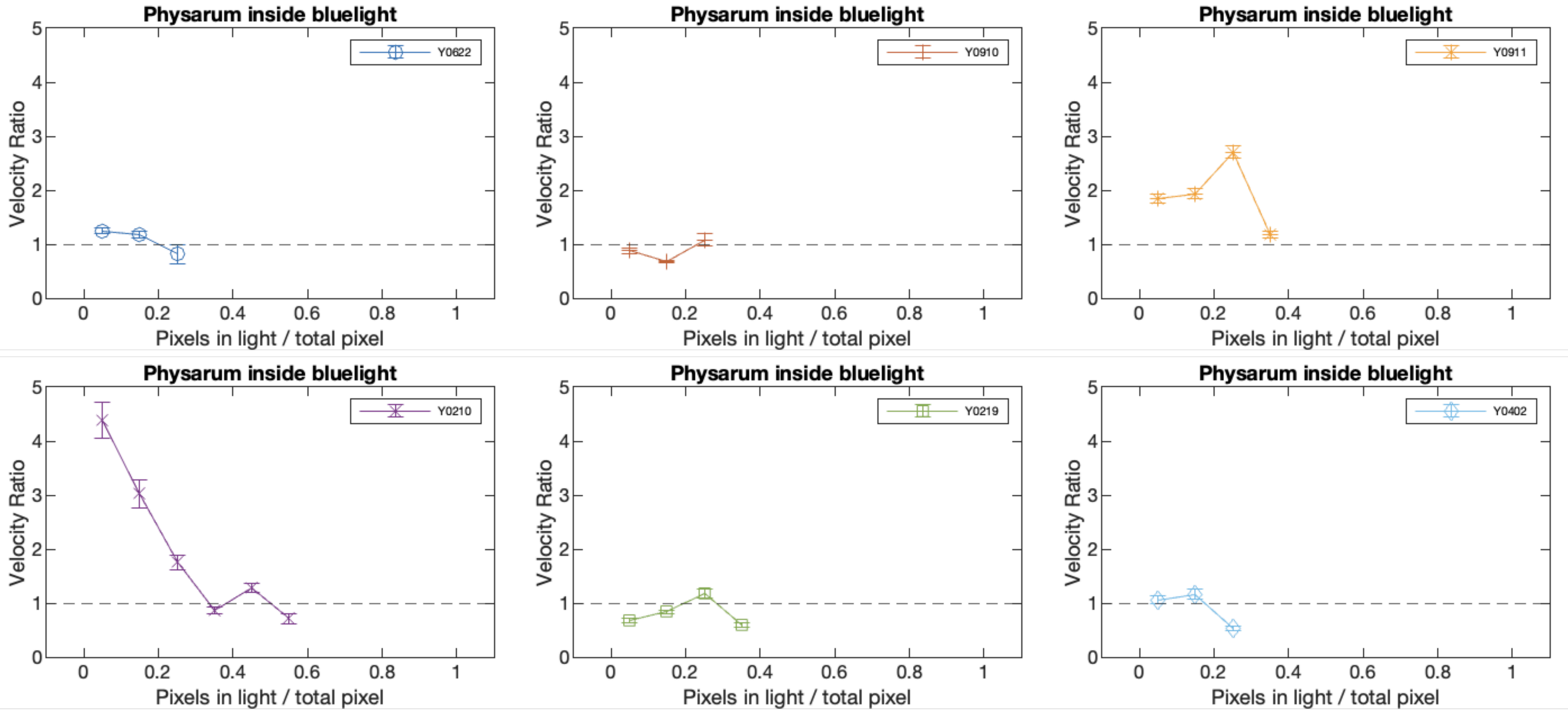


**Supplemental Figure 3.** Change of velocity for the parts exposed to light for individual Y-shaped organisms, plotted against the relative mass of *P. polycephalum* in blue light relative to the whole body. These plots were also plotted in the background of Figure 2 C. Yet 5 out of the 6 specimens showed clear oscillations in velocity during the evacuation response underlining the also on average dynamics shown in Figure 2.
