## Supplementary material for "Network topology enables efficient response to environment in *Physarum polycephalum*": Figure S1

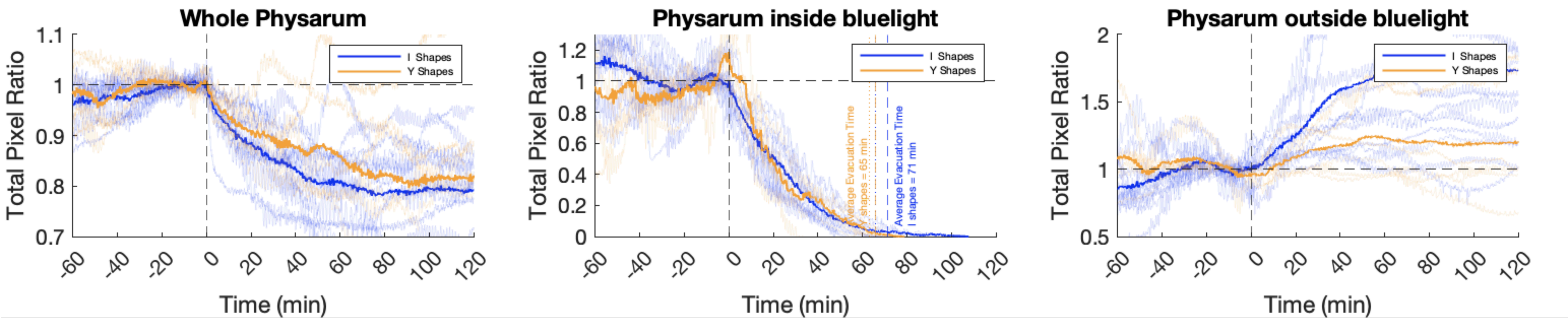


**Supplemental Figure 1.** Figure 1 B-D with x-axis extended to include up to 60 min before light stimulation. Typically, we avoid considering the full first hour of imaging quantitatively as the predominantly first half hour of data acquisition is dominated by *P. Polycephalum's* adaptation to the microscope light. Yet, even without considering the microscopy light adaptation the variation in mass before blue light is applied is considerably less than the mass loss triggered by the blue light.

Note that the distinction between inside and outside blue light is obviously random before blue light is applied rendering higher variability in the total pixel ratio to arise merely be organism foraging dynamics.
